# Stavudine Reduces NLRP3 Inflammasome Activation and Upregulates Aβ-Autophagy

**DOI:** 10.1101/377945

**Authors:** Francesca La Rosa, Marina Saresella, Ivana Marventano, Federica Piancone, Enrico Ripamonti, Chiara Paola Zoia, Elisa Conti, Carlo Ferrarese, Mario Clerici

## Abstract

Alzheimer’s disease (AD) is associated with amyloid-beta (Aβ) deposition and neuroinflammation, possibly driven by activation of the NLRP3 inflammasome. Nucleoside reverse transcriptase inhibitors (NRTI) hamper the assembly of the NLRP3 inflammasome; we analyzed whether stavudine (D4T), a prototypical NRTI, modulates Aβ-mediated inflammasome activation; because neuroinflammation impairs Aβ clearance by phagocytes, phagocytosis and autophagy were examined as well. THP-1-derived macrophages were stimulated in vitro with Aβ_42_ alone or after LPS priming with/without D4T. NLRP3 and TREM2 expression was analyzed by RT-PCR, phagocytosis and ASC-Speck by AmnisFlowSight, NLRP3-produced cytokines by ELISA, authophagy by P-ELISA evaluation of P-ERK and P-AKT. Results showed that IL1β, IL18 and caspase-1 were increased whereas Aβ-phagocytosis and TREM2 were reduced in LPS+Aβ_42_-stimulated cells. D4T reduced NLRP3 assembly as well as IL18 and caspase-1 production, but not IL1β, phagocytosis, and TREM2. P-AKT expression was augmented and P-ERK was reduced by D4T, suggesting a stimulatory effect on autophagy. D4T reduces NLRP3 inflammasome-associated inflammation, possibly restoring autophagy, in an *in vitro* model of AD; it will be interesting to verify its possibly beneficial effects in the clinical scenario.

## INTRODUCTION

Alzheimer’s disease is a neurodegenerative pathology associated with the deposition of extracellular amyloid beta (Aβ) plaques within the brain. Oligomeric Aβ-deposition is held responsible for the activation of microglia, which results in the production of several neurotoxic molecules including inflammatory cytokines, ultimately leading to neurotoxicity, progressive synaptic loss, and cognitive decline (Town *et al*, 2001; Tan *et al*, 2002; Town *et al*, 2005; Townsend *et al* 2005; Simard *et al*, 2006; Town *et al*, 2008; Fiala & Veerhuis, 2010; Feng *et al*, 2011; Heneka *et al*, 2015a).

Inflammation has convincingly been demonstrated to be a key driver of the disease (Tuppo & Arias 2005; Griffin et al, 2006; Cai et al, 2014; Wanga et al, 2017), and recent results suggest that the activation of the nod-like receptor protein 3 (NLRP3) inflammasome is responsible for such inflammation. This is based on a number of observations, thus: 1) the concentration of the inflammasome-derived proinflammatory cytokines interleukin (IL)-1β and IL-18 is increased in AD (Heneka *et al*, 2013; Gold & El Khoury, 2015; Saresella *et al*, 2016; Awad *et al*, 2017); 2) NLRP3-deficiency in the APP/PS1 mouse model of AD decreases neuroinflammation and Aβ accumulation and improves neuronal function (Heneka *et al*, 2013); and 3) higher IL-1β concentrations are detected in individuals with a diagnosis of amnestic mild cognitive impairment (aMCI) that convert into AD (La Rosa *et al*, 2018). Notably, NLRP3 up-regulation is now recognized as a central component in the development of several inflammatory and autoimmune diseases (Lamkanfi *et al*, 2012; Strowig *et al*, 2012; Guo *et al*, 2015).

In AD Aβ accumulation is initially contrasted by the activation of phagocytic cells. In the long run though, these cells become engulfed by Aβ peptides; this leads to the release of cathepsin B, which further stimulates the activation of the NLRP3 inflammasome (Halle *et al*, 2008). This process is contrasted by the action of triggering receptor expressed on myeloid cells 2 (TREM2), a protein that plays a protective role in AD by inhibiting the inflammatory response, enhancing Aβ phagocytosis (Hamerman *et al*, 2006; Casati *et al*, 2018), and promoting microglial functions in response to Aβ deposition (Bouchon *et al*, 2001; Klesney-Tait *et al*, 2006; Neumann & Takahashi, 2007; Bajramovic 2011; Colonna & Wang 2016; Tan *et al*, 2017). The Aβ accumulation-associated phagocytosis defect also jeopardizes the activity of the autophagic flux (Fiala *et al*, 2005; Hui *et al*, 2017). This is important in AD, as alterations of autophagy, a process that mediates lysosomal degradation of proteins, inflammatory cells and organelles, were shown to play a pathogenic role in this disease (Uddin *et al*, 2018). Thus, recent data showed that the degradation of extracellular Aβ by microglia is dependent on autophagic processes, and that autophagy is important for the regulation of Aβ-mediated NLRP3 activation (Rodgers *et al*, 2014; Qian *et al*, 2017).

The mechanism linking autophagy and NLRP3 activation was clarified by observations indicating that authophagy down regulates NLRP3 via the induction of a lysine 63–linked ubiquitination of the NLRP3 adaptor molecule ASC (Shi *et al*, 2012); the autophagosome engulfs these substrates and eliminates them after fusion with the lysosome (Harris *et al*, 2011). Hence, autophagy disrupts multiple steps of inflammasome activation to prevent excessive inflammation (Harris *et al*, 2011; Ferguson TA & Green, 2014; Saitoh *et al*, 2014); The exact intracellular signaling mechanisms involved in the regulation of autophagy are still to be elucidated, but a pivotal role for two MAP-kinases: ERK and AKT, is strongly suspected (Martinez-Lopez *et al*, 2013; Joassard *et al*, 2013). Thus, ERK1 phosphorylation (ERK-P) activates mTOR and, as a consequence, inhibits autophagy; conversely the phosphorylation of AKT (AKT-P) inhibits mTOR (Hay 2005; Peng *et al*, 2010) and activates autophagosomes through chaperone-mediated (LAMP) autophagy (Yang & Klionsky, 2010; Heras-Sandoval *et al*, 2014; Li *et al* 2017). Of note, ERK1/2 phosphorylation was recently shown to inhibit the NLRP3 inflammasome (Mezzasoma *et al*, 2017), and ERK-P was demonstrated to associate with Aβ accumulation in an AD animal model (Jin *et al* 2012).

These observation, together with data implying that inflammasome activation impacts on the risk of developing AD, suggest that targeting inflammation and, in particular, the activation of the NLRP3 inflammasome, could positively modulate Aβ phagocytosis and autophagy, possibly offering a therapeutic opportunity. Recent results showed that nucleoside reverse transcriptase inhibitors, including stavudine (D4T), one of the compounds initially used in the therapy of HIV infection, inhibit the activation of the inflammasome and prevents the transcription of proteins that are part of the inflammasome complex (Kerur *et al*, 2013; Fowler *et al*, 2014;). These results suggest a possible beneficial role of D4T in inflammatory diseases, including AD. We verified this hypothesis in an *in vitro* system using THP-1-derived macrophages.

## RESULTS

### Cellular toxicity of D4T

To determine the optimal dose of D4T to be used in the experiments, two different doses of the drug (50 μM and 100 μM) were added to cell cultures of THP-1-derived macrophages; cell viability was analyzed using the MTT assay. Results showed that, whereas the higher dose of D4T significantly reduced the viability of cells compared to control (medium alone); the lower dose of drug had only a marginally effect on this parameter as >90% of cells were viable at the end of the incubation period (data not shown). Based on these results, the dose of 50 μM D4T was used in all the experiments.

### D4T reduces mRNA expression of NLRP3 proteins in THP-1-derived macrophages

Quantitative PCR analyses were performed in THP-1-derived macrophages that were stimulated with Aβ_42_ alone or with Aβ_42_ after priming with LPS; analyses were performed in the absence/presence of D4T. Stimulation in both conditions: Aβ_42_ alone or Aβ_42_ after priming with LPS, resulted in a significant upregulation of the mRNA expression of all the proteins that are part of the NLRP3 inflammasome complex, reinforcing the idea that Aβ_42_ accumulation results in NLRP3 activation-driven inflammation (Fig 1).

**Figure 1:**
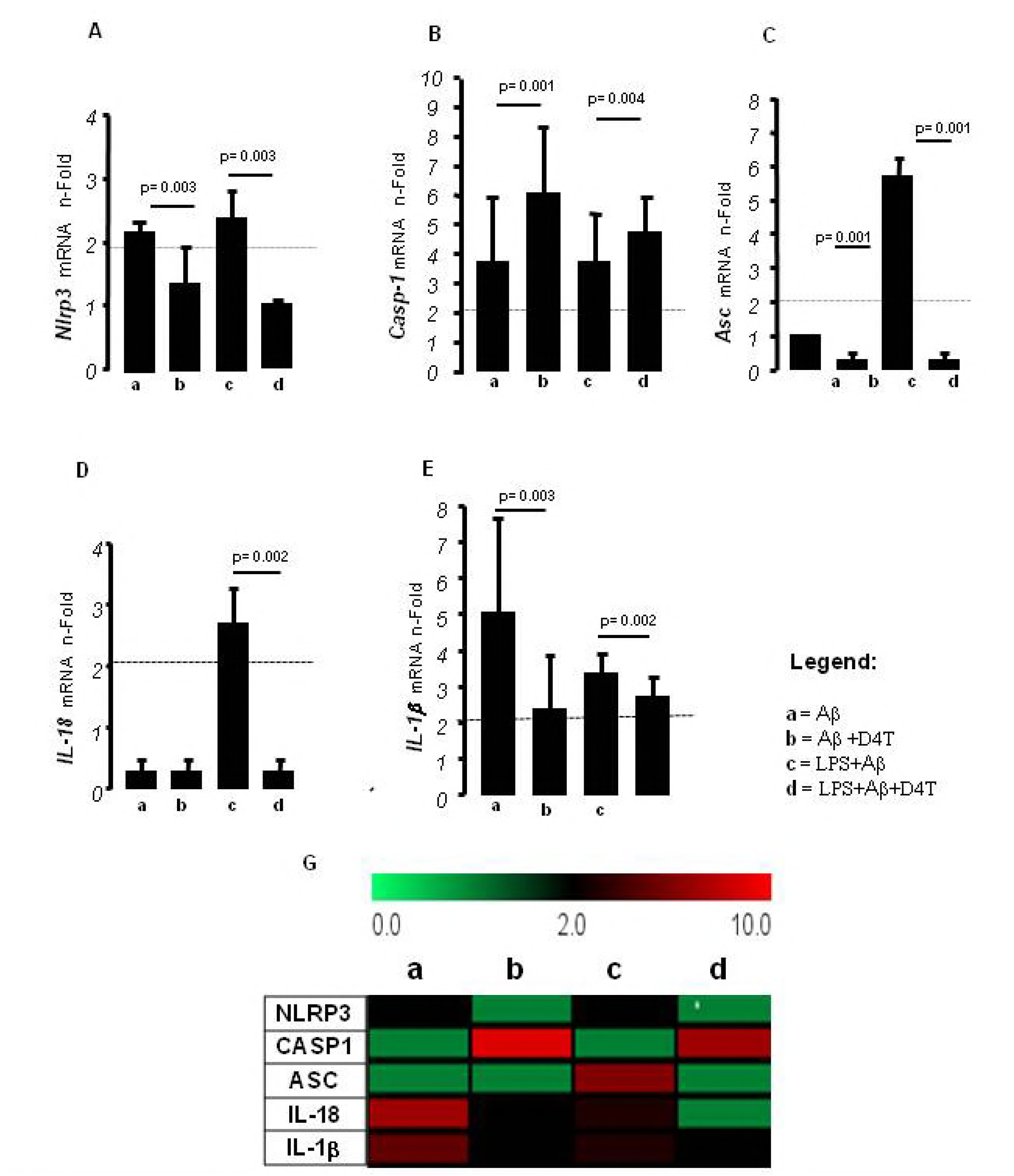
Inflammasome pathway mRNA expression and in stimulated macrophage derived THP-1 cell line; Single real-time PCR results obtained in cells cultured alone with Aβ (10μg/ml) or primed with lipopolysaccharide (LPS) (1μg/ml) and Aβ with or without D4T. Genes of inflammasome proteins: IL-1β, caspase1, IL-18, NLRP3 and ASC are shown in the panels A-F. Gene expression was calculated relative to GAPDH housekeeping gene. The results are indicated as fold-change expression from the unstimulated samples. Summary results, obtained using the TIGR Multi Experiment Viewer (MeV)v4.9, are shown in the bar graphs in panel G.

Notably, D4T was able to significantly reduce the mRNA expression of the sensor (Nlrp3), adaptor (ASC), and effectors (IL-1β and IL-18) proteins that compose the NLRP3 inflammasome (*p* < *0.005* in all cases). In contrast with these results, D4T did not reduce but rather increased mRNA expression of the catalytic NLRP3 inflammasome proteine caspase-1 (Fig 1).

### Effect of D4T on NLRP3-associated production of pro-inflammatory cytokines

The production of the NLRP3 activation-related proinflammatory cytokines IL-1β and IL-18, as well as that of caspase-1, was measured next in THP-1-derived macrophages that were stimulated with Aβ_42_ alone or with Aβ_42_ after priming with LPS in the absence/presence of D4T.

Results showed that the production of all these proteins was up regulated in both Aβ_42_-stimulated and LPS+Aβ_42_-stimulated cells (*p* < *0.05* in all cases), with the highest protein production being observed in LPS+Aβ_42_-stimulated cells.

The addition of D4T significantly reduced IL-18 and caspase-1 production in all the examined experimental conditions (*p* < *0.005* in all cases); the drug significantly down regulated IL-1β production by Aβ_42_–stimulated THP-1-derived macrophages alone as well (*p* < *0.005*), but it had a marginal effect, or, paradoxically, it increased IL-1β production by LPS+Aβ_42_-stimulated cells. These results are shown in Figure 2.

**Figure 2:**
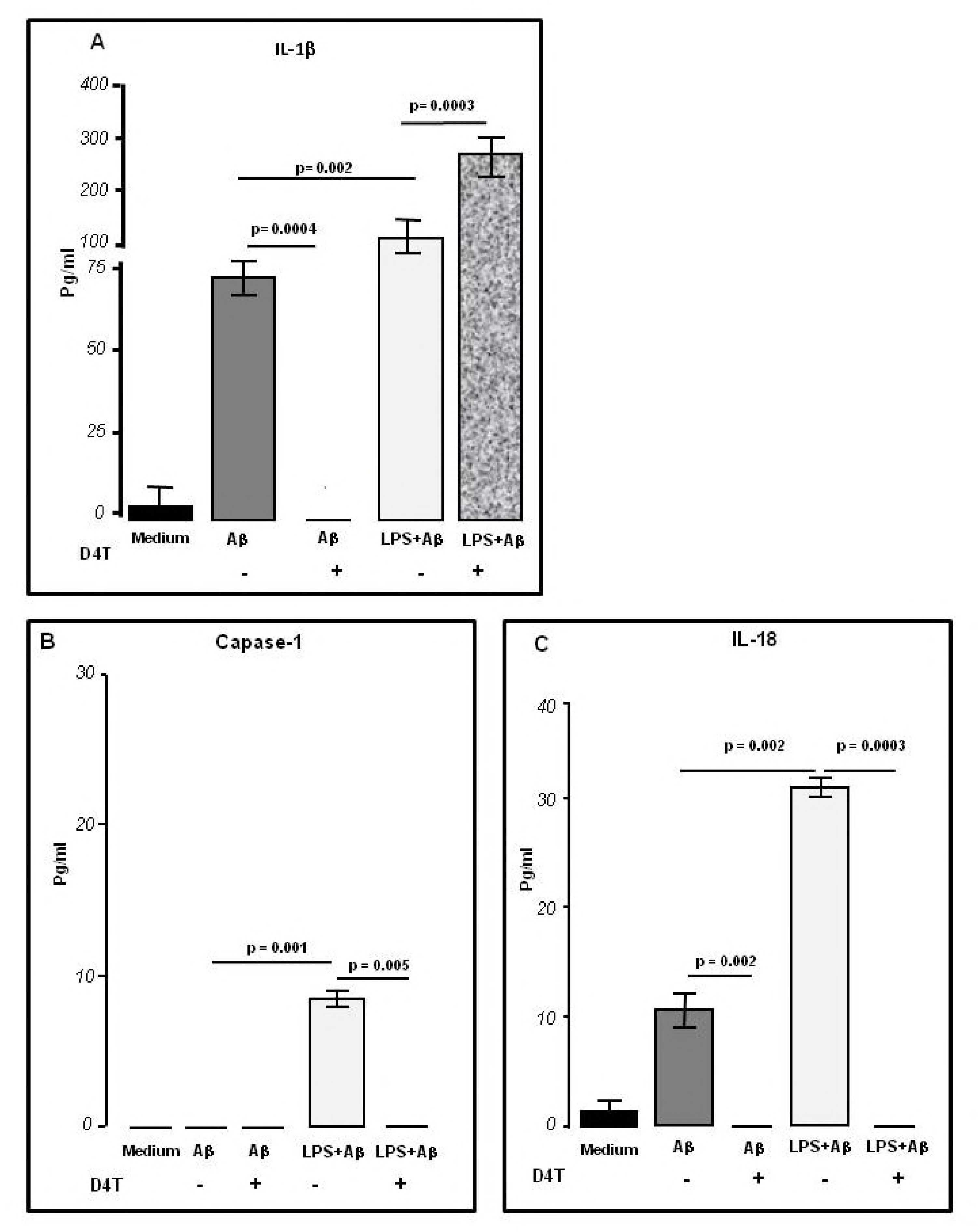
Modulation of NLRP3 inflammasome effector proteins by D4T in stimulated macrophage derived THP-1 cell line; Cytokines IL-1β (panel A), Caspase 1 (panel B), and IL-18 (panel C) production was measured by ELISA on supernatants of THP-1 cells treated with Aβ-alone (10μg/ml) or primed with lipopolysaccharide (LPS) (1μg/ml) and/or with D4T (50μM). Data are representative of three independent experiments and expressed as means±SD. Untreated cells condition was indicated as medium. Statistical significance is shown.

### D4T inhibits NLRP3/ASC-speck inflammasome assembly

The effect of D4T on NLRP3 inflammasome activation was verified next in THP-1-derived macrophages that were stimulated in the same experimental conditions by using the Amnis Flow Sight technology. Representative images are provided in figures 3, panels A-thorough-C. Results confirmed that, as compared to what observed in cells stimulated with Aβ_42_ alone (Fig 3A), LPS+Aβ_42_ stimulation causes a much more extensive assembly of NLRP3 and ASC within large protein complexes (specks), which are the result of inflammasome activation (Fig 3B). Results also showed that D4T prevents the generation of specks, hence impeding the assembly of the NLRP3 inflammasome (Fig 3C). Figure 3D shows overall results that can be summarized as follows: 1) NLRP3 ASC-speck colocalization (assembly of inflammasome) is significantly increased in LPS+Aβ_42_ compared to Aβ_42_ alone-stimulated cells (*p* = *0.0001*); 2) D4T significantly reduces NLRP3/ASC-speck colocalization (assembly of inflammasome) in LPS+Aβ_42_ activated cells (*p* = *0.007*).

**Figure 3:**
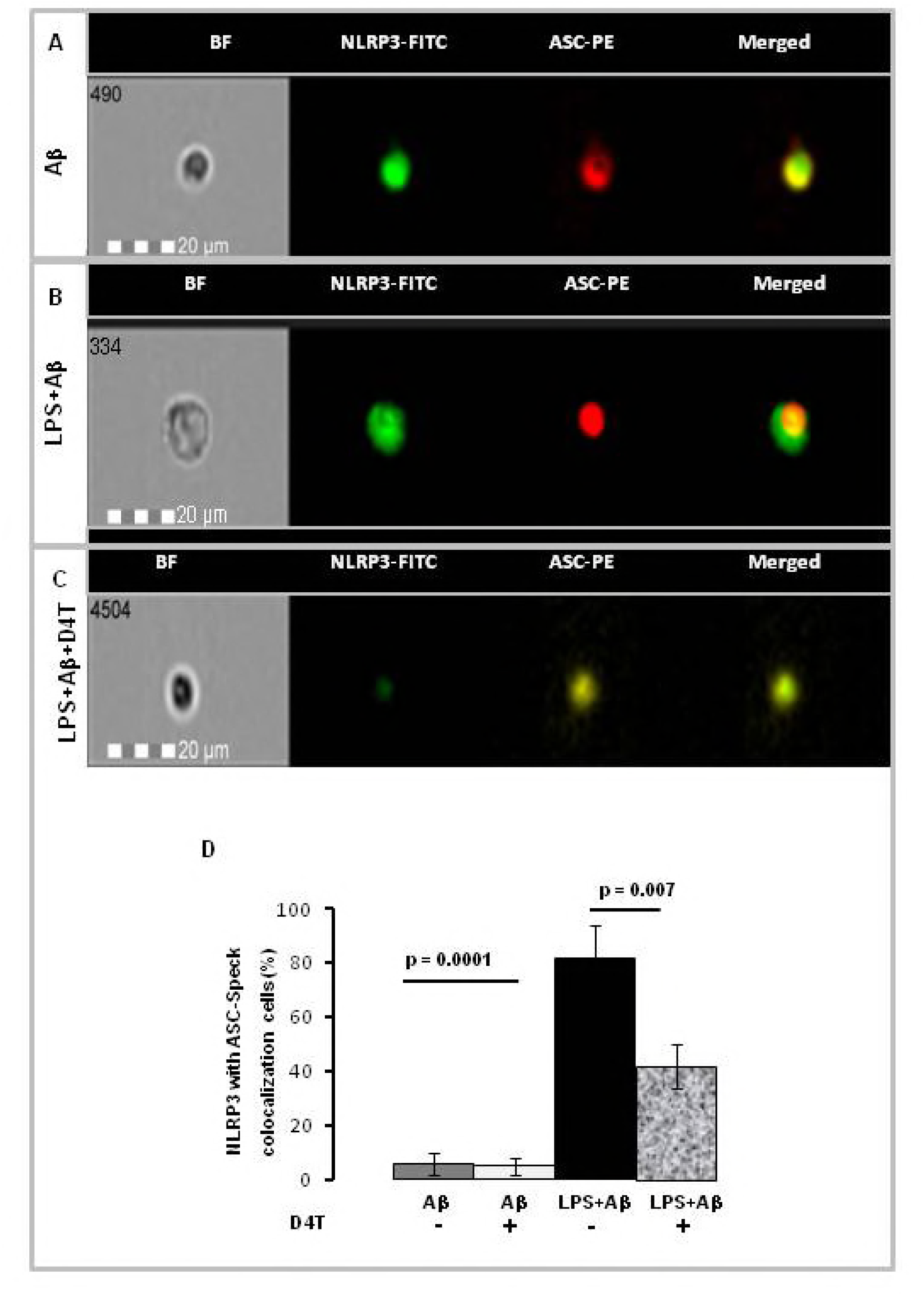
*NLRP3/ASC-speck formation in treated macrophage derived THP-1 cell line*. NLRP3/ASC-speck colocalization was analyzed by Amnis Flow Sight Cytometry. 1*10^6^ cells treated with Aβ alone (10μg/ml) (panel A), or after inflammasome activation with lipopolysaccharide (LPS) (1μg/ml) and Aβ_42_ (LPS+Aβ) (panel B) or after D4T treatment (panel C). In all panels: the first column shows cells in brightfield (BF), second column shows NLRP3-FITC fluorescence, third column shows ASC-PE fluorescence, and fourth column shows florescence of ASC merged with NLRP3. Results were summarized as percentage of positive cells for NLRP3/ASC-speck in LPS+Aβ compared to Aβ and/or with D4T stimulated cells (IDEA software) (panel D). Statistical significance is shown. Scale bar 20 μm.

### D4T modulation of Aβ_42_ phagocytosis by THP-1-derived macrophages

The phagocytic ability of THP-1-derived macrophages that were stimulated with Aβ_42_ alone or with Aβ_42_ after priming with LPS in the absence/presence of D4T was measured next. Results showed that NLRP3 activation was correlated with a significantly reduced ability of these cells to phagocyte Aβ_42_ (*p* ≤ *0.05 in all condition*), suggesting that NLRP3 inflammasome-related inflammation plays a role in the impairment of phagocytosis seen in AD. Results also showed that addition of D4T to cell cultures could not restore the impairment of Aβ_42_ phagocytosis by THP-1-derived macrophages in any of the analyzed experimental conditions (*p* < *0.05 in all condition*). These results are shown in Figure 4.

**Figure 4:**
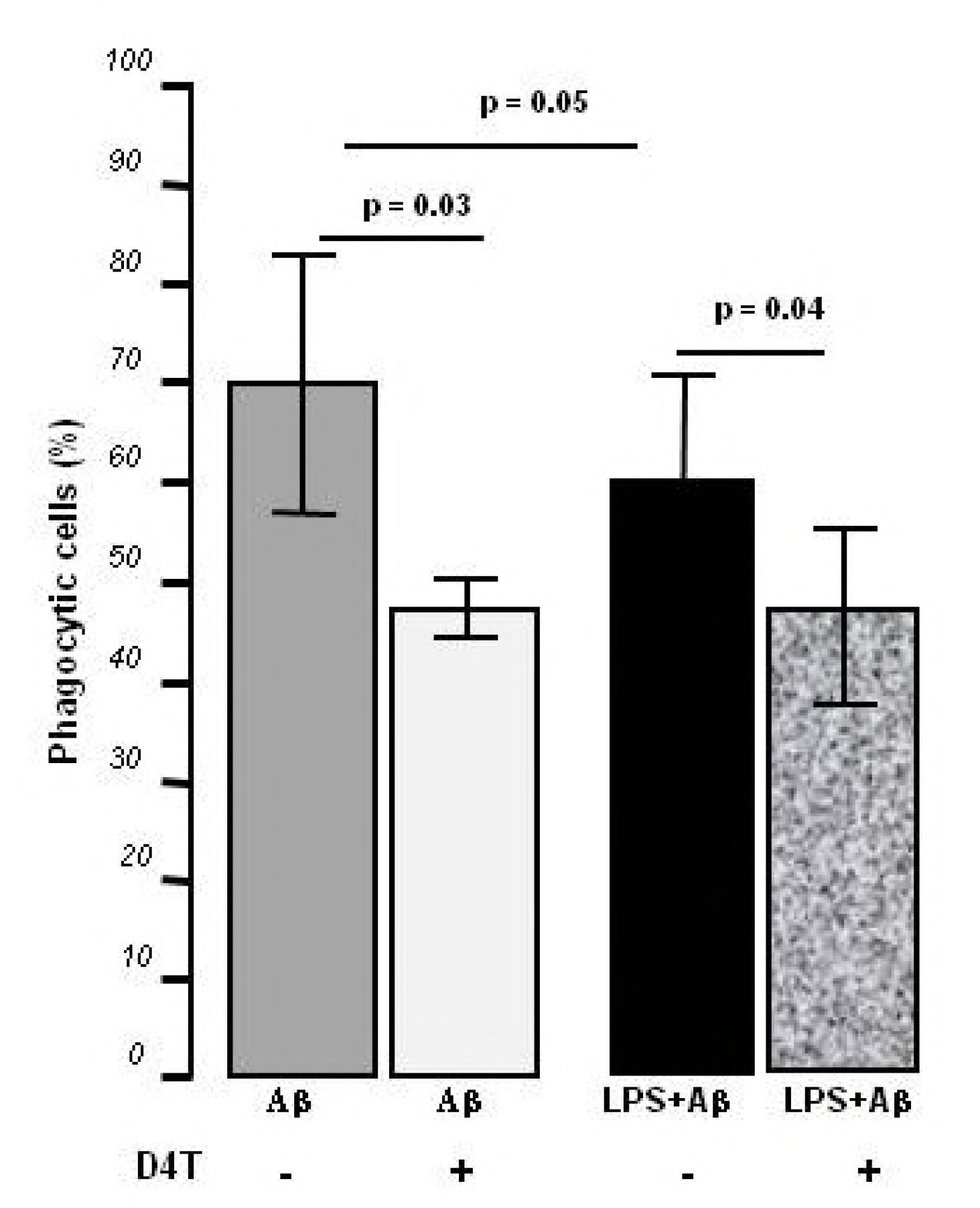
*AMNIS FlowSight analysis of percentage for FAM-* Aβ *positive macrophage derived THP-1 cell line;* The phagocytic ability of THP-1-derived macrophages were measured in all experimental conditions; 1×10^6^ macrophage were treated only with FAM-Aβ or after inflammasome activation with lipopolysaccharide (LPS) (1μg/ml) and FAM-Aβ (LPS+ Aβ) (panel A) and/or with D4T treatment (panel B). Data were shown as mean ± SD of three independent experiments; statistical significance is indicated.

### D4T modulation of TREM-2 mRNA expression

To determine the relationship between NLRP3-activation and Aβ-phagocitosis we also evaluated TREM-2 mRNA expression in all experimental conditions. Results showed a significant reduction of TREM-2 mRNA in LPS +Aβ_42_ compared to Aβ_42_ alone stimulated cells (*p* ≤ *0.001*). When the effect of D4T on TREM-2 expression was analyzed, results indicated that TREM-2 was reduced by D4T in all the examined conditions. (*p* ≤ *0.001*) (Fig 5), further confirming that this compound does not have an effect on phagocytosis.

**Figure 5:**
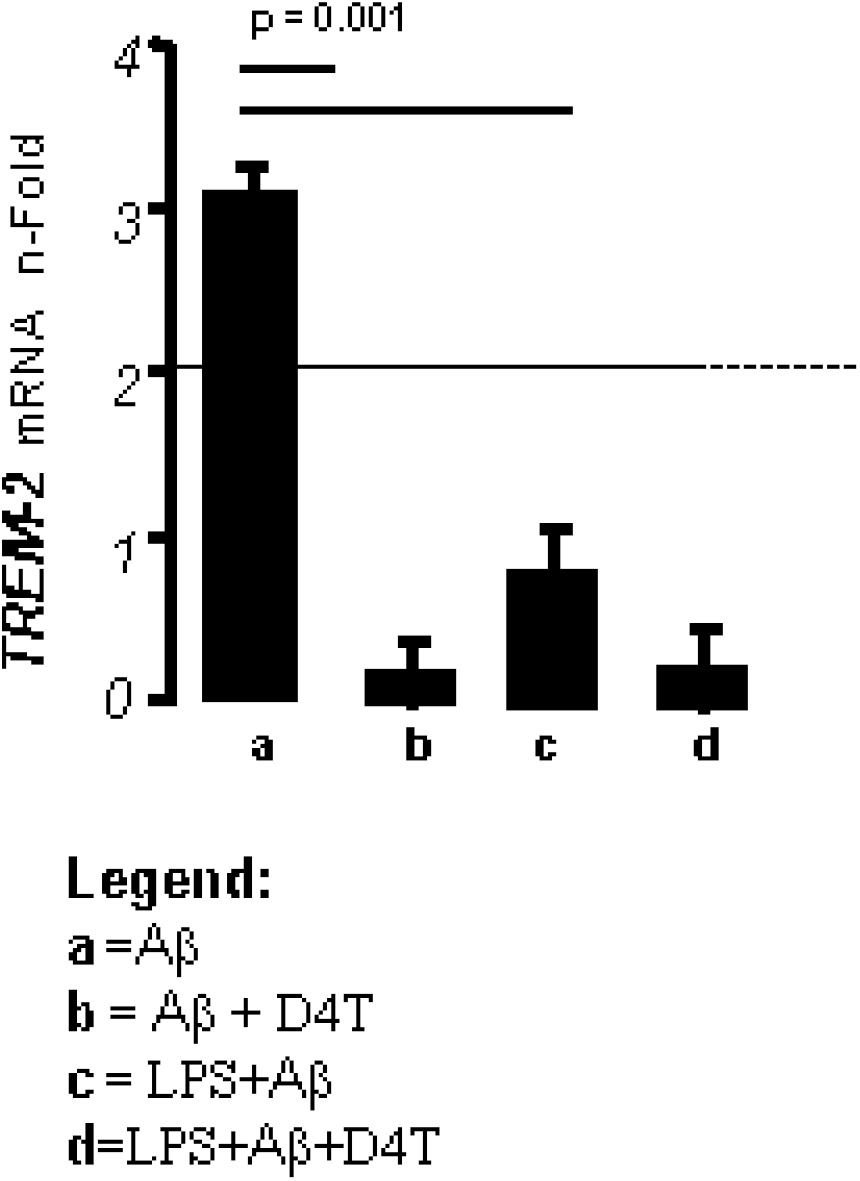
TREM-2 expression in macrophage derived THP-1 cell line; mRNA expression by single real-time PCR; results are indicated as fold-change expression from the unstimulated samples and calculated relative to GAPDH housekeeping gene. Statistical significance is shown.

### D4T-modulation of ERK e AKT-phosphorylation and autophagy

D4T did not modulate Aβ-phagocytosis, but the action of an alternate phagocytic pathway, autophagy, was repeatedly shown to play a primary role in Aβ degradation by the microglia. We thus analyzed the relative activation of these two phagocytic pathways by evaluating the phosphorilation status of ERK and AKT.

Result showed that phosphorylation of these two proteins was significantly modulated by D4T in protein extracts of THP-1-derived macrophages. Thus: 1)D4T resulted in a significant up-regulation of AKT phosphorilation in all conditions (*p* < *0.05 in all conditions*); 2) in the same protein extracts, p-ERK was significantly decreased by D4T in all the examined experimental conditions (p ≤ *0.001* for all comparison) (Fig 6).

**Figure 6:**
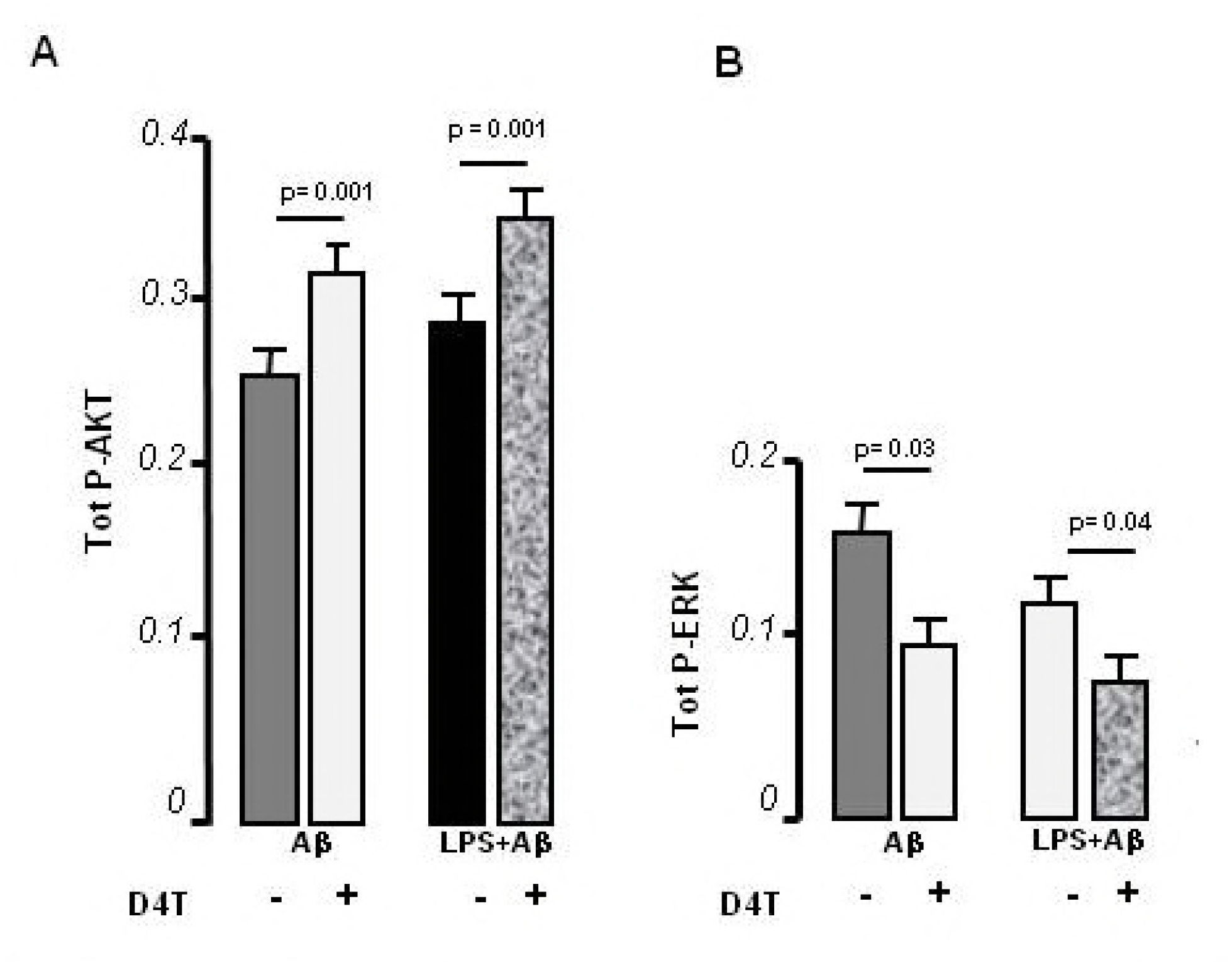
Autophagy kinases modulate NLRP3 in macrophage derived THP-1 cell line; phosphorylation status of ERK (A) and AKT (B) kinases was assessed by phospho(P)-ELISA in cytosol protein extracts of unstimulated cells or lipopolysaccharide (LPS) (1μg/ml) and Aβ stimulated cells and/or with D4T treatment. Data were analyzed by ANOVA and is depicted as mean ± SD of three independent experiments; significant difference from cells untreated is shown.

## DISCUSSION

Inflammasomes are fundamental intracellular structures formed by a number of proteins whose assembly results in inflammation. An exaggerated and persistent activation of the NLRP3 inflammasome, though, has repeatedly been shown to play a pivotal role in autoimmune and inflammatory diseases including AD. The involvement of NLRP3 inflammasome in AD, in particular, is supported by results obtained both in patients and in animal models, including the observations that the concentration of IL-1β and IL-18, prototypical NLRP3-produced proteins, is increased in AD (Heneka *et al*, 2013; Gold & El Khoury, 2015; Saresella *et al*, 2016; Awad *et al*, 2017) and that NLRP3-deficiency in the APP/PS1 mouse model of AD decreases neuroinflammation and Aβ accumulation and improves neuronal function (Heneka *et al*, 2013). Further support to the involvement of the NLRP3 inflammasome in the pathogenesis of AD was offered by results showing that Aβ induces the processing of pro-IL-1β into mature IL-1β in the microglia via activation of NLRP3 inflammasome (Parajuli *et al*, 2013), and that NLRP3 inflammasome deficiency favors the differentiation of microglia cells into the M2 phenothype (anti-inflammatory) (Hu *et al*, 2015; Dempsey *et al*, 2017). Neuroinflammation also impacts on the ability of the phagocytes to eliminate Aβ peptides, and in AD it is well known that an impairment in the ability of phagocytes to catabolize Aβ peptides leads to the engulfment and the functional paralysis of these cells, favoring the accumulation of Aβ and its deposition in plaques. We used an *in vitro* system to analyze the involvement of NLRP3 inflammasome activation in Aβ-phagocytosis and to verify whether the dampening of inflammasome activation would result in a stimulation of autophagy. To this end we used Stavudine (D4T) an antiviral designed to target HIV reverse trascriptase, that was recently shown to be endowed with the ability of down modulating NLRP3 inflammasome activation. Data herein, obtained using an *in vitro* model of AD and methods that allow single cell direct visualization by merging flow cytometry and high-resolution microscopy, show that the NLRP3 inflammasome activation is directly correlated with the impairment in phagocytosis that characterizes AD. We also demonstrate that D4T, an antiviral that has been widely used in the therapy of HIV infection, greatly reduces NLRP3 inflammasome activation and, whereas it does not have an effect on phagocytosis, it could significantly upregulate Aβ authophagy by macrophages.

D4T down regulated the expression of all NLRP3 proteins mRNA with the exception of caspase-1 This is possibly the result of a moderate activation of mRNA caspase-1 by LPS priming, or, alternatively, it could be driven by the spontaneous release of endogenous ATP (Netea *et* al, 2009). Similarly, the drug reduced the production of IL-18 but not of IL1β; this discrepancy can be justified by the observation that, whereas IL-18 is a purely NLRP3 activation-derived cytokine, IL1β can be secreted by monocytes independently of classical inflammasome stimuli (Gaidt *et al*, 2016). Aβ clearance is a complex, multifactorial process, requiring the collaboration of various systems and cell types, including microglia, macrophages and peripheral monocytes (Zuroff *et al*, 2017). Notably, whereas it is still unclear whether Aβ accumulation is a cause or consequence of disease, mounting evidences have shown that increased cerebral Aβ burden is the earliest pathologic event in AD, supporting the idea that Aβ accumulation plays a principal role in this disease. Clearance of Aβ is believed to be hampered as a consequence of inflammation. Our data showing a significant reduction of Aβ -phagocytosis in THP-1 derived macrophages that were preactivated with LPS (inflammatory condition) compared to those cultured with Aβ alone (noninflammatory condition) support this idea. Notably, thou, whereas D4T could greatly reduce NLRP3 inflammasome activation, this compound did not have any effect of phagocytosis in the in *in vitro* experimental model we used. The lack of effect of D4T on phagocytosis was further reinforced by the observation that the expression of TREM2, a receptor that is expressed on microglial/macrophages cells (Jones *et al*, 2014) and acts as a sensor for Aβ clearance, was not modulated either by this compound. This discrepancy could be explained in different ways: 1) Aβ phagocytosis is independent from NLRP3 activation; 2) other, yet unknown mechanisms impair phagocytosis even when NLRP3 activation is impeded; 3) our system does not represent what goes on *in vivo*. An alternate and more interesting explanation stems from a number of recent results suggesting that successful Aβ clearance in AD is mediated not by phagocytosis but, rather, by autophagy: a process that mediates lysosomal degradation of proteins, inflammatory cells and organelles.

Autophagy and NLRP3 inflammasome activation have indeed been linked by the observation that authophagy down regulates NLRP3 *via* the induction of a lysine 63–linked ubiquitination of ASC and, on the other hand, autophagy inhibition exacerbates inflammasome activity and disease in models of influenza infection, and autoimmune conditions including IBD (Nakahira *et al*, 2011; Lupfer *et al* 2013; Ravindran *et al*, 2016).

Autophagy is a complex metabolic mechanism that includes a number of pathways, the best characterized of which are microautophagy, which is mediated by the phosphorylation of AKT, and LAMP chaperone-mediated autophagy (Kiffin *et al*, 2004; Vernon & Tang, 2013). We observed that, in concomitance with its dampening effect on NLRP3 activation, D4T resulted in a significant increase of phosphorylated AKT. AKT phosphorylation modulates mTOR signaling pathway and LAMP chaperon-mediated autophagy. These results confirm, at least in the *in vitro* system we used, that D4T could have a beneficial effect on stimulating autophagy-mediated Aβ clearing possibly as a consequence of its ability to reduce NLRP3 inflammasome activation. These data could be important in the light of observations in the animal model of AD showing that, in autophagy deficient mice, accumulation of Aβ is seen in brain cells and results in neurodegeneration and memory impairment (Nilsson *et al*, 2013). Besides increasing AKT phosphorylation, D4T also significantly reduced the phosphorylation of another protein: ERK. Notably, whereas Ras-ERK signaling induces Aβ and τ hyperphosphorylation, which is characteristically observed in AD brains, p-ERK-p down regulation prevents τ and Aβ phosphorylation as well as neuronal cell cycle entry (Kirouac *et al*, 2017). This complex link was further reinforced by recent results showing that ERK phosphorylation is reduced in the presence of low Aβ concentrations (Kirouac *et al*, 2017).

Stavudine (D4T), an antiviral, has been used since the mid-80s in millions of HIV-infected individuals (Fowler *et al*, 2014); this compound was shown to prevent caspase-1 activation and to be efficient in mouse models of geographic atrophy, choroid neovascularization and graft-versus-host disease (Fowler *et al*, 2014). Results herein indicating that D4T down regulates NLRP3 inflammasome activation and stimulates Aβ autophagy in an *in vitro* model of Alzheimer’s disease seem to warrant the investigation of its possible use in the clinical scenario.

## MATERIALS AND METHODS

### Cells

THP-1 human monocytes (IZSLER, Istituto Zooprofilattico Sperimentale della Lombardia e Dell’Emilia Romagna, IT) were grown in RPMI 1640 supplemented with 10% FBS, 2mM L-glutamine, and 1% penicillin (medium)(Invitrogen Ltd, Paisley, UK). To differentiate these cells into macrophages, monocytes were seeded in 6-well plates at a density of 1.0×10^6^ cells/well in medium that contained 50ηM of phorbol 12-myristate 13-acetate (PMA)(Sigma-Aldrich, St. Louis, MO) and incubated for 12 h at 37°C in 5% CO2; cells were then resuspended in serum-free medium.

### Cell culture

THP-1-derived macrophages were cultured with medium alone or incubated with: 1) Aβ_42_ (10μg/ml)(Anaspec, Fremont, California, USA) for one hour, or 2) Aβ_42_ after a 23 hours priming with Lypopolissacaride (LPS) (1μg/ml)(Sigma-Aldrich) in the absence/presence of D4T (50μM)(Sigma-Aldrich)(Lachlan R. Gray et al, 2013). Alexa Fluor 488 (FAM)-labeled Aβ_42_ was used in the phagocitosis assayes and in ASC-speck detection; non-labeled Aβ_42_ was used for gene-expression and protein quantification.

### Cellular toxicity

Viablity of THP-1-derived macrophages was determined using the MTT 3-(4,5-dimethylthiazol-2,5-diphenyltetrazolium bromide) (Sigma-Aldrich) assay, as previously described (Mossmann, 1983).

### RNA extraction and reverse transcription

RNA was extracted from 1×10^6^ THP-1-derived macrophages in unstimulated or stimulated conditions (see above) in the absence/presence of D4T and reverse-transcribed into first-strand cDNA (Biasin *et al*, 2010). Real-time PCR cDNA quantification was performed by real-time PCR as previously described (Biasin *et al*, 2007).

### NLRP3-Inflammasome and Trem2 gene expression

NLRP3, ASC, Caspase-1, IL-1β, IL-18 (Qiagen, Hilden, Germany) and TREM-2 (Sigma-Aldrich) expression was evaluated by RT-PCR. Results were expressed as ΔΔCt (where Ct is the cycle threshold) and are presented as the ratio between the target gene and the GAPDH housekeeping mRNA.

### NLRP3-downstream inflammasome protein quantification by ELISA

Proinflammatory cytokines were analyzed in supernatants of THP-1-derived macrophages in unstimulated or stimulated conditions (see above) in the absence/presence of D4T. Caspase-1, IL-1β and IL-18 concentration was analyzed by sandwich immunoassays according to the manufacturer’s recommendations (Quantikine Immunoassay; R&D Systems, Minneapolis, MN, USA). A plate reader (Sunrise, Tecan, Mannedorf, Switzerland) was used and optical densities (OD) were determined at 450/620 nm. All the experiments were performed in triplicates. Sensitivity (S) and Assay Range (AR) were as follows: S: IL-1β=1pg/ml; Caspase-1= 1.24 pg/ml; IL-18= 12.5 pg/ml. AR: IL-1β 3.9 −250 pg/ml; Caspase-1= 6.3 - 400 pg/ml; IL-18= 25.6-1000 pg/ml.

### AMNIS FlowSight analysis

Aβ-FAM-phagocytosis, ASC-speck formation and NLRP3-complex assembly were analyzed by FlowSight (Amnis Corporation, Seattle, WA). 1×106 in THP-1-derived macrophages stimulated as described above. Cellswere fixed with 100 μl of PFA (1%) (BDH, UK), permeabilized with 100 μl of Saponine (0.1%) (Life Science VWR, Lutterworth, Leicestershire, LE) and stained with FITC-anti human NLRP3 (Clone #768319, isotype Rat IgG2a, R&D Systems,) and PE-anti human ASC (clone HASC-71, isotype mouse IgG1, Biolegend, San Diego, CA, USA) for 1 h at room temperature. Cells were then washed with PBS, centrifuged at 1,500 rpm for 10 min and resuspended in 50 μl of PBS; results were analyzed by IDEAS analysis software (Amnis Corporation, Seattle, WA, USA).

The FlowSight is an imaging flow cytometer that together merges flow cytometry and high-resolution microscopy. It is equipped with two lasers operating at 488 and 642 nm, two camera and twelve standard detection channels. It simultaneously produce side scatter (darkfield) image, one or two transmitted light (brightfield) images, and up to ten channels of fluorescence imagery of every cell. FlowSight using the InspireTM system, acquires 2000 cells/second and operates with a 1 Dm pixel size (~20X magnification) allowing visualization of fluorescence from the membrane, cytoplasm, or nucleus; the IDEAS image analysis software allows quantification of the fluorescence at different cellular localizations.

Phagocytosis assay were performed by internalization feature utilizing a mask representing the whole cell, defined by the brightfield (BF) image, and an internal mask defined by eroding the whole cell mask in order to eliminate the fluorescent signal coming from Aβ_42_-FAM attached to the cell surface, thus measuring only the internalized part. The internalization feature was first used to calculate the ratio of the intensity of FAM (Aβ_42_ signal) inside the cell/ total FAM intensity outside the cell. Higher internalization scores indicate a greater concentration of Aβ_42_ FAM, inside the cell (Supplementary Figure).

ASC-speck and NLRP3-inflammasome assembly analyses were performed using the same mask of internalization feature, differentiating for ASC diffuse or spot (speck) fluorescence inside of cells and its co-localization with the NLRP3 inflammasome protein.

### Phagocytosis assay

1×10^6^ THP-1-derived macrophages were cultured with Alexa Fluor 488 (FAM)-labeled Aβ_42_ for 1h, in unstimulated or stimulated conditions (see above) in the absence/presence of D4T. Medium containing 0.05% Trypsin-EDTA (COD. ECB3052D, Euroclone, Milan, IT) was then added for 10 minutes at 37°C in 5% CO_2_; to block trypsin, cells were then resuspended in RPMI 1640 supplemented with 10% FBS and centrifuged at 1500 rpm for 10 min. Pellet was fixed with 0.1% paraformaldehyde (PFA) for 10 min, washed and resuspended in 50μl PBS. Supernatants were collected and stored at −80°C for cytokine measurement by ELISA.

### ERK1/2 and AKT quantification: total-andphospho-ELISA

Phosphorilated ERK1/2 and AKT in lysates of THP-1-derived macrophages were analyzed using Immunoassay Kits (phosphor-ELISA kits, BioSource International, Inc). Because total levels of ERK1/2 and AKT are independent of phosphorylation status, total ELISA kits (BioSource) were used to normalize the phosphorylated ERK1/2 and AKT content of the samples.

Protein cell extraction was performed in Cell Extraction Buffer (Biosource), containing 1mM PMSF, protease and phosphatase inhibitor cocktail (Sigma-Aldrich)(1:200 and 1:100), for 30min, on ice. Lysates were then centrifuged at 12000g for 10 minutes at 4°C. Different dilutions of samples were tested for each phosphorylated or total protein detection. Protein absorbance was determined at 450nm (BioRad); concentrations were calculated comparing absorbance to the specific standard curve values for phosphorylated ERK and AKT, and expressed with respect to each specific protein kinase total status.

### Statistical analysis

Experiments were repeated at least three times with triplicate of each condition. Firstly we performed a parametric analysis of variance (one-way ANOVA) to evaluate phagocytosis and cytokine production. Repeated measures ANOVA and Tukey post-test were performed for kinases analyses. Results of ANOVA models are shown as means and SD (standard deviations). Post hoc comparisons were run using t tests with Tuckey’s HSD (honestly significant difference) procedure. Data analysis was performed using the MedCalc and R statistical packages. Results were considered to be statistically significant if surviving the p <0.05 threshold.

## Acknowledgements

This work was supported by 2017–2018 Ricerca Corrente (Italian Ministry of Health).

## Author contributions

MC came up with the idea. FLR, MS and MC designed the experimental procedure. FLR, IM and FP project performed the experiments. CPZ, EC and CF contributes reagents and useful insights. ER performed statistical analyses. FLR, MS and MC wrote and edited the manuscript.

## The paper explained

ProblemIn early phase of dementia, deposition of beta amyloid (Aβ) plaques induce neuroinflammatory processes and the activation of immune system which include the activation of microglia and the recruitment of peripheral monocytes in the AD brain. In particular, the inflammasome has been implicated in several chronic inflammatory and autoimmune diseases and recent data *(Halle 2008, Heneka 2013, Saresella 2016)* showed that NLRP3, an inflammasome component, is activated in AD. NLRP3 activation leads to production of the proinflammatory cytokines IL-1ß, IL-18 and caspase-1 resulting in inflammatory milieu that reduce Aβ-clereance and down-regulation of TREM2 *(Thornton 2017)* a receptor protein that allows microglia to phagocyte Aβ.

## Results

Recent data show that nucleoside reverse transcriptase inhibitors (NRTI) are endowed with an intrinsic anti-inflammatory activity as they inhibit the assembly of the inflammasome. Given these data, we hypothesize that the administration of NRTI and in particular Stavudine (an oral compound that has been used for years in HIV infection) blocking NLRP3 assembly, could prevent caspase-1 activation and, as a consequence, IL-18 release; this could blocking inflammasome, or could polarize monocytes to an anti-inflammatory phenotype, restoring phagocytosis functions. We propose that this drug could be interesting tool in the treatment of AD.

## Impact

An in-depth analysis of the possible beneficial effects of component that suppress inflammasome NLRP3 activation in AD (NRTI-stavudine)..

## Supplementary data

Aβ_42_-phagocytosys detection by FlowSight specifically shows single cell events that were identified using a dot plot of brightfield (BF) Aspect Ratio *versus* BF Area Aspect Ratio is a feature value calculated by dividing the height of the cell by the width. On the FlowSight single cell events tend to have an aspect ratio between 0.7 and 1.0. On the basis of macrophages a second plot, using Aβ-internalization score (IS) was generated to identify the percentage of THP-1-derived macrophages were positive for Aβ-FAM relative to the negative control A positive value of IS corresponds to a cell with mostly Aβ-internalized, represents the ratio of fluorescence intensity inside the cell to the total fluorescence intensity of the cell and identifies the phagocytic capacity of the cells.

<u>*Supplementary figure: Determination of Aβ phagocytosis by macrophage derived THP-1 cell line;*</u>

Representative images capture by Amnis Flow Sight Cytometry of cells untreated (negative control) or stimulated with Alexa Fluor 488-labeled Aβ (FAM-Aβ); In panel A are shown gated macrophages image (left) based on Area and Aspect Ratio: area of the cell were identified as number of pixels and converted to μm^2^ (1 pixel = 0.25 μm^2^); image of percentage of FAM-Aβ positive macrophages (positive control) is shown (right). In panel B is shown brightfield (BF) image (left) of negative cell for Alexa Fluor 488 fluorescence (right). In panel C: first column shows BF image of macrophages treated with FAM-Aβ (a), second column shows image related to FAM-Aβ fluorescence (b), third column shows merged fluorescence with BF (c). Internalization score (IS) calculated by IDEA software is shown (d). Scale bar 20 μm.

